# Viral fitness predicts the magnitude and direction of perturbations in the infected host transcriptome

**DOI:** 10.1101/206789

**Authors:** Héctor Cervera, Silvia Ambrós, Guillermo P. Bernet, Guillermo Rodrigo, Santiago F. Elena

## Abstract

Determining the fitness of viral genotypes has become a standard practice in virology as it is essential to evaluate their evolutionary potential. Darwinian fitness, defined as the advantage of a given genotype with respect to a reference one, is a mesoscopic property that captures into a single figure differences in performance at every stage of viral infection. But to which extent viral fitness results from particular molecular interactions with host factors and regulatory networks during infection? Can we identify host genes, and then functional classes, whose expression depends on viral fitness? Here, we compared the transcriptomes of tobacco plants infected with seven genotypes of tobacco etch potyvirus (TEV) that differ in fitness. We found that the larger the fitness differences among genotypes, the more dissimilar the transcriptomic profiles are. Consistently, two different mutations, one in the viral RNA polymerase and another in the viral suppressor of RNA silencing, that led to close fitness values, also resulted in significantly similar gene expression profiles. Moreover, we identified host genes whose expression showed a significant correlation, positive or negative, with TEV fitness. Over-expression of genes with positive correlation activates hormone-and RNA silencing-mediated pathways of plant defense. By contrast, under-expression of genes negatively correlated reduces metabolism, growth, and development. Overall, these results reveal the high information content of viral fitness, and suggest its potential use to predict differences in genomic profiles of infected hosts.

---

Fitness is a complex parameter often used by evolutionary biologists and ecologists to quantitatively describe the reproductive ability and evolutionary potential of an organism into a particular environment^1,2^. Despite this apparently simple definition, measuring fitness is difficult and most studies only measure one or more fitness components (*e.g*., survival to maturity, fecundity, number of mates, or number of offspring produced) as proxies to total fitness^1,2^. In the field of virology, it has become standard to measure fitness by growth-competition experiments in mixed infections with a reference strain^3,4^. With this experimental set up, fitness is just the relative ability of a viral strain to produce stable infectious progeny in a given host (cell type, organ, individual, or species) when the available resources have to be shared with a competitor^5^. Regardless its limitations, at least, this approach provides a metric for ranking viral strains according to their performance in a particular environment/host. Such a fitness measure has been pivotal for quantitatively understanding many virus evolution processes: the effect of genetic bottlenecks and accumulation of deleterious mutations in RNA virus populations^6–8^, the rates and dynamics of adaptive evolution into novel hosts^9,^, the pleiotropic cost of host range expansion^10–12^, the cost of genome complexity^13,14^, the cost of antiviral escape mutations^15–17^, the topography of adaptive fitness landscapes^18–20^, and the role of robustness in virus evolution^21–23^.

But differences in viral fitness should also matter in genome wide studies seeking to understand the mode of action of the viruses (*i.e*., the precise way they interact with their hosts). Even though it has been argued that an integrative systems biology approach to viral pathogenesis would result in a better understanding of pathogenesis and in the identification of common targets for different viruses, therefore serving as a guide to a more rational design of therapeutic drugs^24–30^, pioneering studies have ignored the high genetic variability of viruses and subsequent differences in fitness and in mode of action. Experimental evidences support that even single nucleotide substitutions have significant effects on viral fitness, regardless they are synonymous or nonsynonymous, or they affect coding or non-coding genomic regions^31–37^. A common trend among all these studies is that, whenever fitness is evaluated in the standard host, the distribution of mutational effects is highly skewed towards deleterious effects, with a large fraction of mutations being lethal. Furthermore, increasing evidences suggest that the shape and location of the distribution of fitness effects depends on the host species, being the difference among two hosts (*e.g.*, a tested host and the natural one) greater as their genetic divergence increases^12,20,38,39^. Together, all these observations suggest that the fitness of a given viral genotype depends not only on its own genetic background, but also on the host where fitness is evaluated. Arguably, differences in viral fitness reflect differences in the virus-host interaction, with respect to a reference strain. As the cellular parasites they are, viruses need to kidnap all sort of cellular factors and resources, reprogram gene expression patterns into their own benefit, and block and interfere with cellular defenses. And all these processes take place embedded into the host complex network of intertwined interactions and regulations. Interacting in suboptimal ways with any of the elements of the host network, due to the accumulation of mutations on the viral genome that affect the functional viral components (*i.e*, RNAs and proteins), may have profound effects in the progression of a successful infection and thereof in viral fitness; inefficient interactions may result in attenuated or even abortive infections.

However, little is known on how viral fitness informs about the underlying changes occurring in host gene expression and protein function at a genome wide scale. In this work, we have investigated the potential association between viral fitness and host transcriptional regulation upon infection as a first step into this direction. To address this, we characterized the transcriptomic profiles of *Nicotiana tabacum* plants inoculated with a collection of genotypes of *Tobacco etch virus* (TEV; genus *Potyvirus*, family Potyviridae) that differ in their fitness in this natural host (Fig. 1). Analyses of expression data allowed us to characterize differential gene expression upon infection with each TEV genotype, as well as to identify sets of candidate genes whose expressions positively or negatively correlate with the magnitude of TEV fitness. Moreover, differences in expression for representative genes from these two categories were experimentally validated by an alternative method.

**Figure 1.**
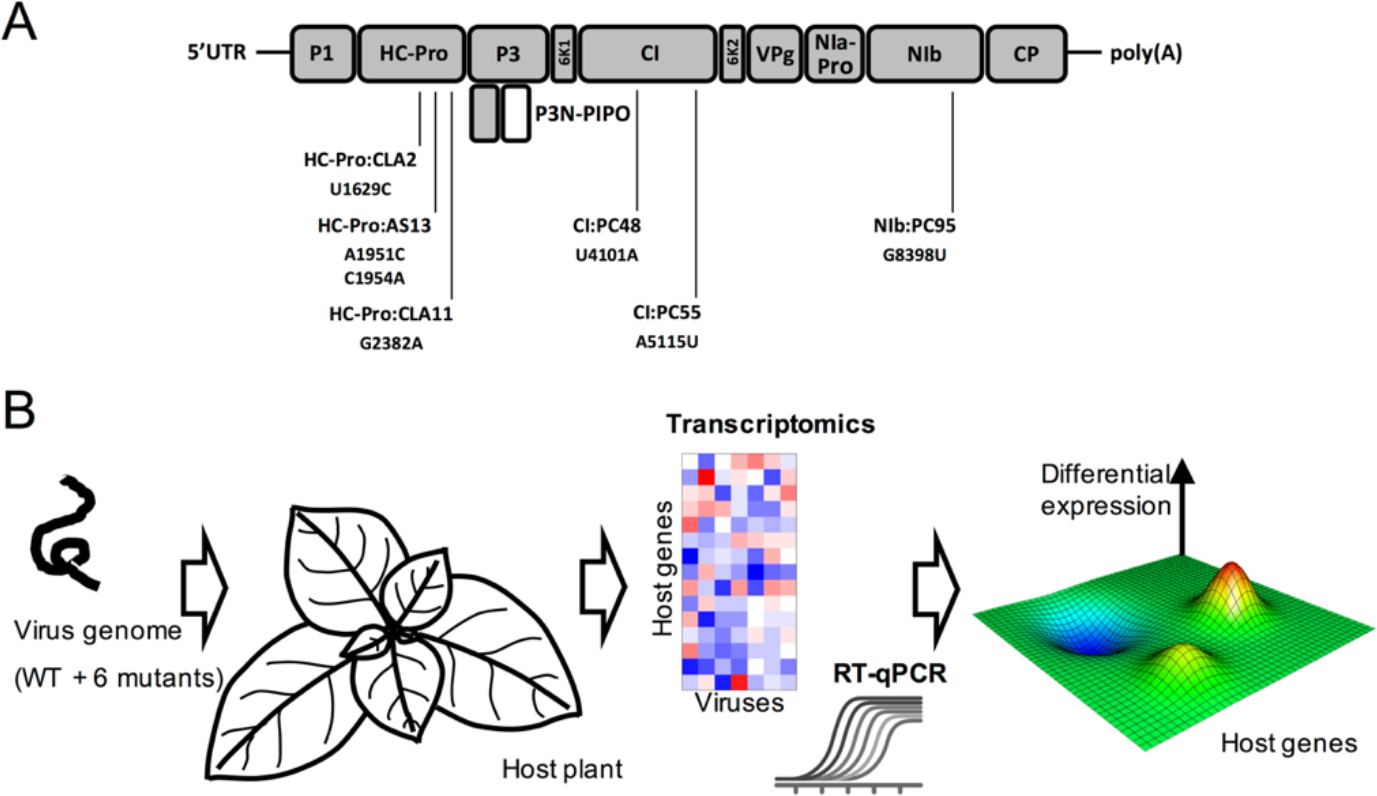
Summary of the experiment. **(A)** Schematic representation of the TEV genome together with the location of the mutations used in this study. **(B)** Schematic representation of the systems biology approach used in this study. A common host plant (*N. tabacum*) was infected with seven different genotypes of TEV differing in their fitness. Viral fitness was evaluated by RT-qPCR, whilst the effect of viruses on the transcriptome of the plants was evaluated by using microarrays.

## Results

### Differences in viral fitness and host symptomatology

Table 1 shows relevant information about the seven TEV genotypes used for this study. The mutant genotypes differ from the wild-type (WT) TEV in a rather limited number of nonsynonymous mutations (1 or 2). However, their fitness values and the severity of symptoms induced differ widely. Significant differences existed among the fitness values of the seven viral genotypes chosen for this study (Fig. 2A; GLM likelihood ratio test: *χ*^2^ = 373.006, 6 df, *P* < 0.001) and among plants inoculated with the same viral genotype (*χ*^2^ = 2927.885, 14 df, *P* < 0.001). As a measure of the quality of data, the percentage of total variance for relative fitness explained by true genetic differences among genotypes was 70.56%, whilst differences among plants inoculated with the same viral genotype accounted for 29.16% of the observed variance. The remaining 0.28% of the observed variability was assignable to error measurements. A Bonferroni *post hoc* test classifies the seven genotypes into five groups (labeled as *a* to *f* in Fig. 2A). Interestingly, the three genotypes with the lowest fitness values (AS13, CLA2 and CLA11) contain mutations in the multifunctional protein HC-Pro, whose most relevant role during infection is to serve as suppressor of the RNA-silencing (VSR) defense^40^. The two genotypes with mutations in the CI protein (PC55 and PC48), a helicase also involved in cell-to-cell movement^40^, have the mildest deleterious fitness effects. Mutant PC95, which contains a point mutation in the replicase NIb protein^40^ occupies an intermediate position in the fitness scale (Fig. 1A).

**Table 1.** Genotypes used in this study and some of their relevant properties.

| **Mutant**<br>**Label** | **Mutated**<br>**Protein** | **Nucleotide**<br>**changed<sup>1</sup>** | **Amino acid**<br>**changed<sup>2</sup>** | **Phenotype** | **Symptoms** |
| --- | --- | --- | --- | --- | --- |
| Wild-type |  |  |  |  | Etch |
| AS13 | HC-Pro | A1951C/A1954C | E299A/D300A | Hyposuppressor | Chlorotic spots |
| CLA2 | HC-Pro | U1629C | V192A | Hyposuppressor | Etch |
| CLA11 | HC-Pro | G2382A | E443K | Hypersuppressor | Mild etching |
| PC48 | CI | U4101A | H156Q | High fitness | Etch |
| PC55 | CI | A5115U | K494N | Low fitness | Mild etching |
| PC95 | NIb | G8398U | A473S | Low fitness | Etch |

<sup>1</sup>Position in the genome<sup>2</sup>Position in the mature peptide

**Figure 2.**
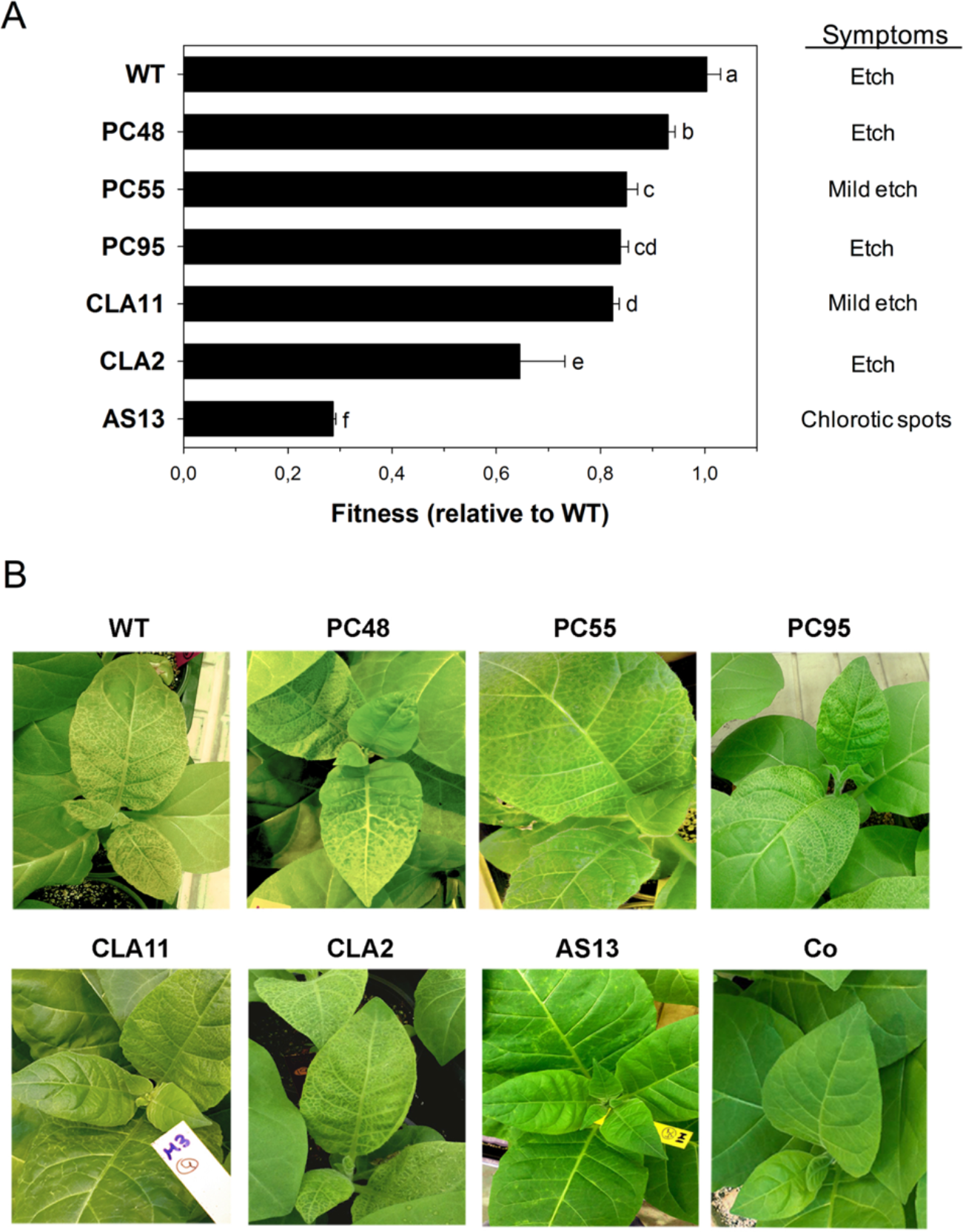
Fitness of the distinct TEV genotypes and symptomatology of the plants. **(A)** Fitness values estimated for each one of the TEV genotypes included in the study in the natural host plant *N. tabacum*. Error bars represent ±1 SEM. Letters (*a* - *f*) next to the bars define the homogenous groups identified using a *post hoc* Bonferroni test. **(B)** Representative plant pictures of the symptomatology induced by each one of the TEV genotypes.

Fig. 2B illustrates the differences in symptoms induced by each one of the seven genotypes. Symptoms ranged from the asymptomatic infection or local chlorotic spots characteristic of mutant AS13, the mild etching of mutant CLA11 and the severe etching induced by the WT and the other mutants. No correlation exists between virus fitness and symptoms, a finding previously reported for this experimental system^32^.

### Differences in viral fitness are associated with differences in the magnitude of the perturbation over the host transcriptome

First, we sought to test whether differences in TEV fitness might be associated with differences in the gene expression profiles of the infected plants. We hypothesized that viral fitness results from a particular interaction between virus and host factors, assuming that the outcome of infection of a WT virus in its natural host results from an *optimal* (from the virus perspective) modulation of the host’s gene expression profile. As viral fitness in the host is reduced, interactions are less optimal and, consequently, the gene expression profile of the plant will be more and more different from that resulting from the successful infection with the WT genotype. To test this hypothesis, we infected *N. tabacum* plants with each one of the seven TEV genotypes described above. Eight days post-inoculation (dpi) symptomatic tissues were collected for all mutants except for the very low fitness mutant AS13, for which tissues were collected 15 dpi because the delay in symptoms appearance and severity (Fig. 2B). Total RNAs were extracted, its quality verified, concentration normalized and used to hybridize *N. tabacum* Gene Expression 4×44K Microarrays (Agilent). Slides were handled as described in the Methods section; intensity signals were normalized using tools in BABELOMICS^41^. Normalized expression data are contained in Supplementary Dataset S1. Fig. 3A shows the clustering (unweighted average distance method; UPGMA) of average expression data for those genes that significantly changed expression (±2-fold) among plants infected with the seven viral genotypes (1-way ANOVAs with FDR correction; overall *P* < 0.05) relative to the mock-inoculated plants.

Regarding individual genes, two major clusters can be distinguished, one corresponding to the over-expression of genes related to stress response and a second one corresponding to the under-expression of genes involved with metabolism and plant development. To further explore the similarity in the perturbation induced by each viral genotype into the plants’ transcriptome, we computed all pairwise Pearson product-moment correlation coefficients (*r*) between the mean expression values for all genes in the microarray. Then, these correlations were used as a measure of similarity to build a UPGMA dendrogram. The rationale for this analysis is as follows: the more correlated two expression profiles are, the more similar the effects induced in infected plants. When comparing expression profiles from a pair of infected plants, a significant correlation may indicate that genes that changed expression relative to the mock-inoculated plants, are *exactly the same* in both samples, showing a similar expression pattern. If this is the case, the correlation coefficient is expected to be high. Conversely, if genes with differential expression do not match in the two samples being compared, then the correlation will be lower. Fig. 3B shows both the heat-map of the correlation coefficients and the resulting dendrogram. Three clusters result from this analysis (Fig. 3B). The first cluster is constituted by the three viral genotypes with the higher fitness values, *i.e*, WT, PC55 and PC48. Genotypes of intermediate fitness CLA11, PC95 and CLA2 constitute a second cluster. Finally, plants infected with AS13 show the most dissimilar gene expression profile. The heat-map is shown with viral genotypes ordered according to the UPGMA clustering. Obviously, it is symmetric, with the diagonal corresponding to the correlation between profiles from plants infected with the same viral genotype, thus being *r* ≈ 1. Correlations decreased as the distance in the cladogram increases. Within clusters, *r* > 0.85, whilst between clusters the correlations ranged 0.65 < *r* < 0.75, except for plants infected with AS13, whose similarity with other infected plants was always *r* < 0.65.

**Figure 3.**
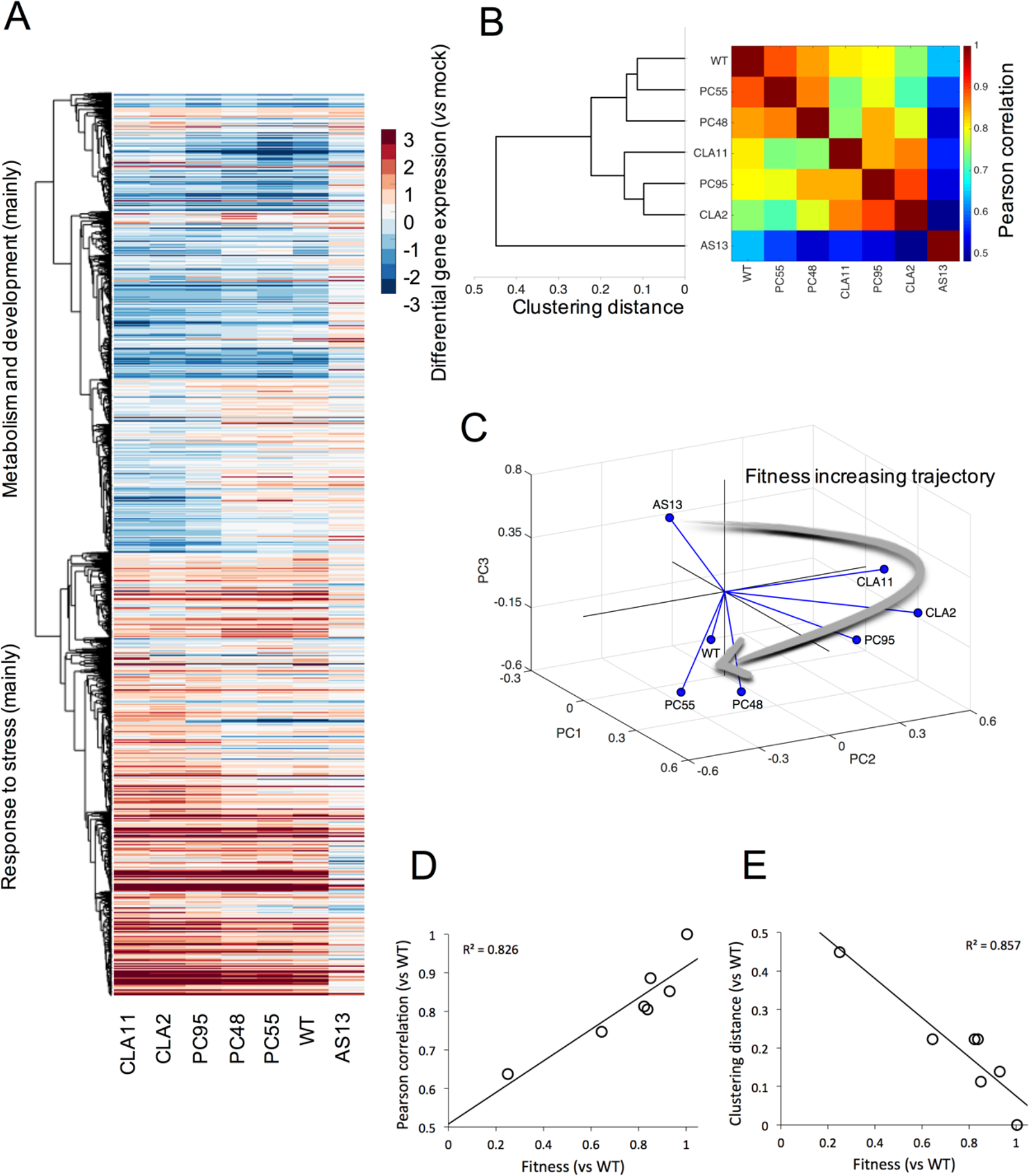
Analyses of the transcriptomic data from tobacco plants infected with each TEV genotype. **(A)** Heat-map representation of gene expression values (infected *vs.* mock-inoculated control plants) for all viral genotypes. Only genes that show significant differences among all infections (ANOVA with FDR, adjusted *P* < 0.05) are included in the heat-map. Hierarchical clustering of genes done with the unweighted average distance method (UPGMA) by using the correlations between all pairs of mean profiles as distance metric. Genes down-regulated (in blue) mainly correspond to metabolic and developmental processes, while genes up-regulated (in red) mainly correspond to stress responses. **(B)** TEV genotypes clustered (UPGMA) according to the similarity of the mean expression profiles of plants infected with each one of them. The heat-map represents the value of the Pearson’s correlation coefficient between pairs of mean profiles. **(C)** Representation of the three major principal components resulting from a PCA of the data shown in panel (B). The three first PCs explain up to 93% of the total observed variance. Lines link each genotype with the centroid of the 3D space. The gray arrow represents a putative trajectory of increasing viral fitness. (D) Association between viral fitness and the magnitude of the perturbation (*vs.* mock-inoculated control plants) both relative to WT (*P* = 0.005). **(E)** Association between viral fitness and the distance of each genotype to theWT (*P* = 0.003), from the dendrogram shown in panel (B).

Next, to further investigate the similarity between expression profiles of plants infected with different TEV genotypes, we performed principal components analysis (PCA) of all the gene expression data. The percentage of total observed variance explained by the first three PCAs was ~93% (the first principal component, *pc*_1_, itself explained up to 81%). Fig. 3C shows the distribution of values in the space defined by the three first principal components. Results are equivalent to those obtained with the two previous clustering methods, as genotypes are classified into three groups. WT, PC55 and PC48 are closer in the space and characterized by positive values of *pc*_1_ but negative values of the second (*pc*_2_) and third (*pc*_3_) components. CLA11, PC95 and CLA2 form a second group, with positive values of *pc*_1_ and *pc*_2_ but negative values of *pc*_3_. As before, AS13 effect on host’ transcriptome is clearly different, and has negative values of *pc*_1_ and *pc*_2_ but positive of *pc*_3_. Interestingly, Fig. 3C shows that genotypes are located in this principal component space following a trajectory of increasing fitness values (indicated by the grey arrow in Fig. 3C). Along this trajectory, *pc*’s switch of sign but in different ways. This transition suggests that the over-or under-expression of a set of genes is associated with particular levels of viral fitness: low fitness AS13 is characterized by a positive *pc*_3_ and a negative *pc*_1_ while high fitness viruses are characterized by the opposite sign. That is, over-or under-expressed genes are not progressively accumulated as long as viral fitness changes. These genes will be evaluated in the following sections.

Following from our working hypothesis, if the WT virus has evolved to optimize its interaction with the host, it is logical that small departures in viral fitness will be associated with small deviations between the transcriptomes of plants infected with the WT virus and with viruses whose fitness are close to the WT. Conversely, the less similar fitness between the WT and mutant viruses, the more dissimilar would be the transcriptional profiles of infected plants. To test this prediction, we have explored (*i*) the correlation between the similarity of transcriptional profiles of plants infected with the WT TEV and with each mutant (again using Pearson’s *r*) and fitness and (*ii*) the correlation between the distance from WT in the cladogram shown in Fig. 3B and fitness. The results of these analyses are shown in Figs. 3D and 3E. As expected, both correlations were significant (*r* = 0.826, *P* = 0.005 and *r* = −0.857, *P* = 0.003, respectively; in both cases 5 d.f.) and of the expected sign.

### The number of altered genes depends on viral fitness

Fig. 4A (data contained in Supplementary Dataset S2) shows the number of differentially expressed genes (DEG), both up-and down-expressed, relative to the transcriptome of mock-inoculated plants. The number of down-expressed DEGs ranges between 531 (for AS13) and 2809 (for CLA11), while in the case of up-expressed DEGs the range is similar: from 781 (AS13) to 2696 (CLA11). Fig. 4B illustrates the number of DEGs in common between all pairs of transcriptomes from infected plants. The heat-map shows a pattern of modularity, with three well defined modules. The first module contains the three viruses with highest fitness (WT, PC48 and PC55), the second module contains the three viruses with intermediate fitness (PC95, CLA11 and CLA2), and the very low fitness genotype AS13 is the only member of the third module. The number of shared DEGs within each of these modules is > 75% of total. The number of shared DEGs between modules drops below 60%.

**Figure 4.**
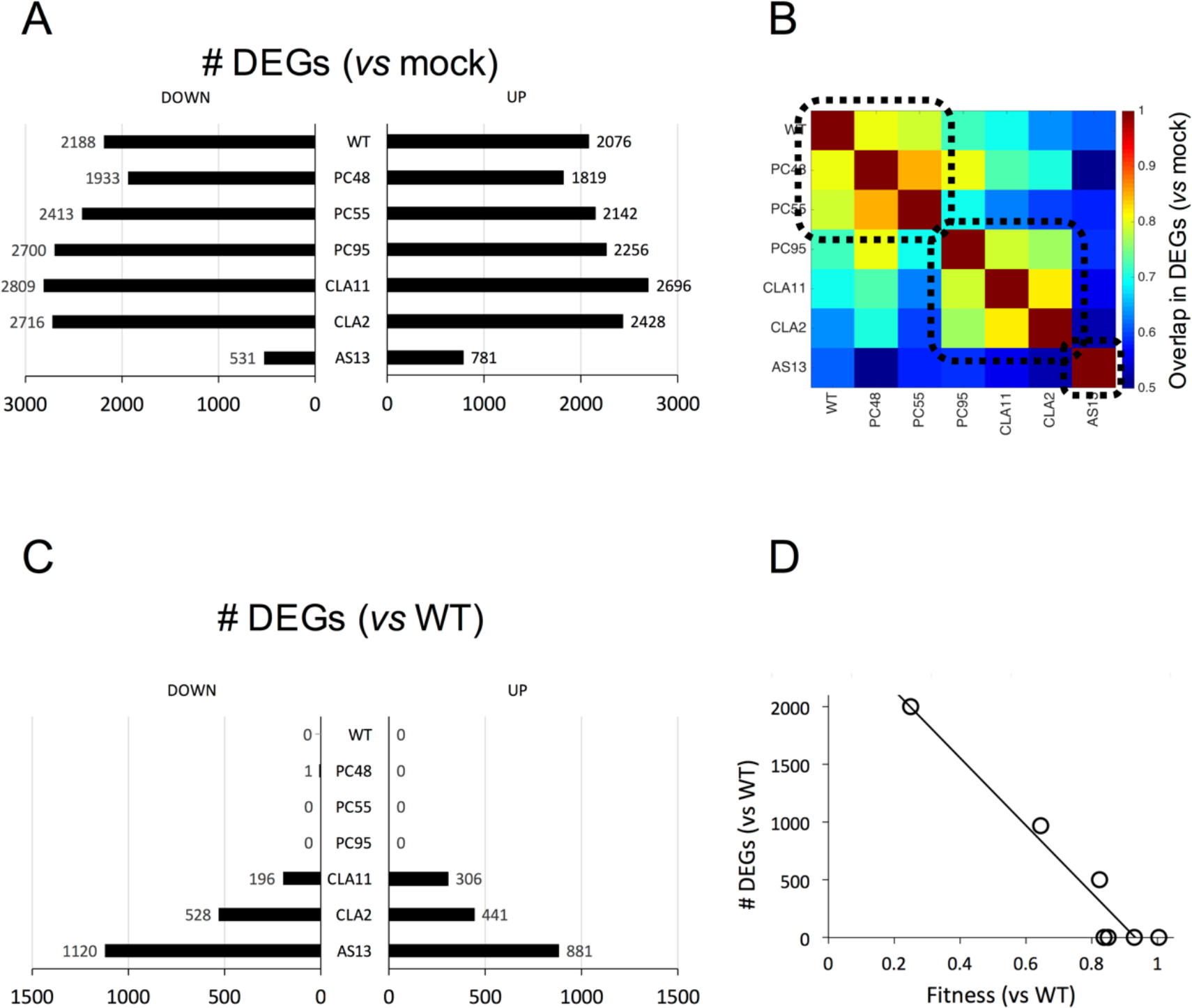
Comparative analysis of differential gene expression in plants infected with the distinct TEV genotypes. **(A)** Distribution of the number of DEGs (up-and down-regulated *vs.* mock-inoculated control plants) for each TEV genotype. **(B)** Heat-map of the degree of overlapping between two lists of DEGs. Three different groups are highlighted (dashed squares). **(C)** Distribution of the number of DEGs (up-and down-regulated *vs.* plants infected with the WT TEV) for each TEV genotype. **(D)** Correlation plot between the total number of DEGs relative to WT TEV infection and the fitness of each TEV genotype (*P* < 0.001).

Next, following the same rationale than in the previous section, we sought to determine whether the number of DEGs also depends on the difference in fitness between WT and the mutant TEV genotypes. In this case, we hypothesized that the overlap in the lists of DEGs must be similar for WT and viruses of equivalent fitness (*e.g*., PC48 or PC55), whereas the magnitude of the overlap between DEG lists would decrease as differences in fitness exist. Fig. 4C shows the counts of DEGs that are differentially expressed (for up-and down-expressed genes) between WT and the other six viral genotypes. As expected, PC48, PC55 and PC95 alter the same genes, though in a different magnitude (as shown in the previous section). The number of genes that are not in common with WT increases from CLA11 (502), CLA2 (969) and AS13 (2001). A highly significant negative correlation exists between the number of differences in DEGs list (adding up the number of up-and down-expressed DEGs in Fig. 4C) and differences in viral fitness (Fig. 4D: *r* = −0.937, 5 d.f., *P* < 0.001).

Of particular interest following all results presented above is the similarity between PC95, a mutant of the replicase *NIb* gene, and CLA11, a mutant of the VSR *HC-Pro* gene. These two mutations led to close fitness values (Fig. 2A), but also resulted in significantly similar gene expression profiles (Fig. 4B). At first sight, one may argue that their impact in transcriptomic profiles should be different since these mutations affect virus proteins that are functionally unrelated. However, our results suggest that the effects on the overall virus-host interaction of each mutant are canalized in the same way. This clearly exemplifies that viral fitness, despite its incompleteness, is a figure that contains high information about the virus-host interaction.

### Viral fitness conditions the functional categories that are altered

Lists of genes are somehow hard to interpret and functional analyses provide a good tool to cluster genes into groups with related functions. To this end, we performed an analysis of enriched functional categories (GO terms) for each viral genotype. Fig. 5 illustrates the way that plants infected with each one of the seven TEV genotypes differ in the functional categories significantly over-represented relative to the mock-inoculated plants. Fig. 5 shows the GO terms in planes ordered from the highest to the lowest viral fitness. In red, we represent categories that are significantly enriched by up-expressed DEGs, in blue categories that are significantly enriched by down-expressed DEGs, and in pink categories enriched in both types of DEGs; the surface of each circle is proportional to the number of DEGs included in each category. The upper plane shows the functional categories altered in plants infected with WT TEV, with *metabolic process* (GO:0008152) containing the largest number and *photosynthesis* (GO:0015979) the smallest one. *Regulation of response to biotic stimulus* (GO:0002831), *defense response* (GO:006952), *immune system process* (GO:0002376), *protein modification process* (GO:0036211), *hormone-mediated signaling* (GO:0009755), and *cell death* (GO:008219) are all enriched in up-expressed DEGs, while *photosynthesis*, *lipid metabolic process* (GO:0006629), and *regulation of nitrogen compound metabolic process* (GO:0051171) are categories significantly enriched in down-expressed DEGs. More wisely defined categories such as *metabolic process*, unspecific *response to stress* (GO:0006950), *response to stimulus* (GO:0050896), *response to abiotic stimulus* (GO:0009628), *localization* (GO:0051641), or *transport* (GO:0008150) are enriched in both types of DEGs. Therefore, overall speaking, genes involved in different aspects of plant defense pathways and response to infection are up-expressed, while genes involved in metabolism and photosynthesis are down-expressed.

**Figure 5.**
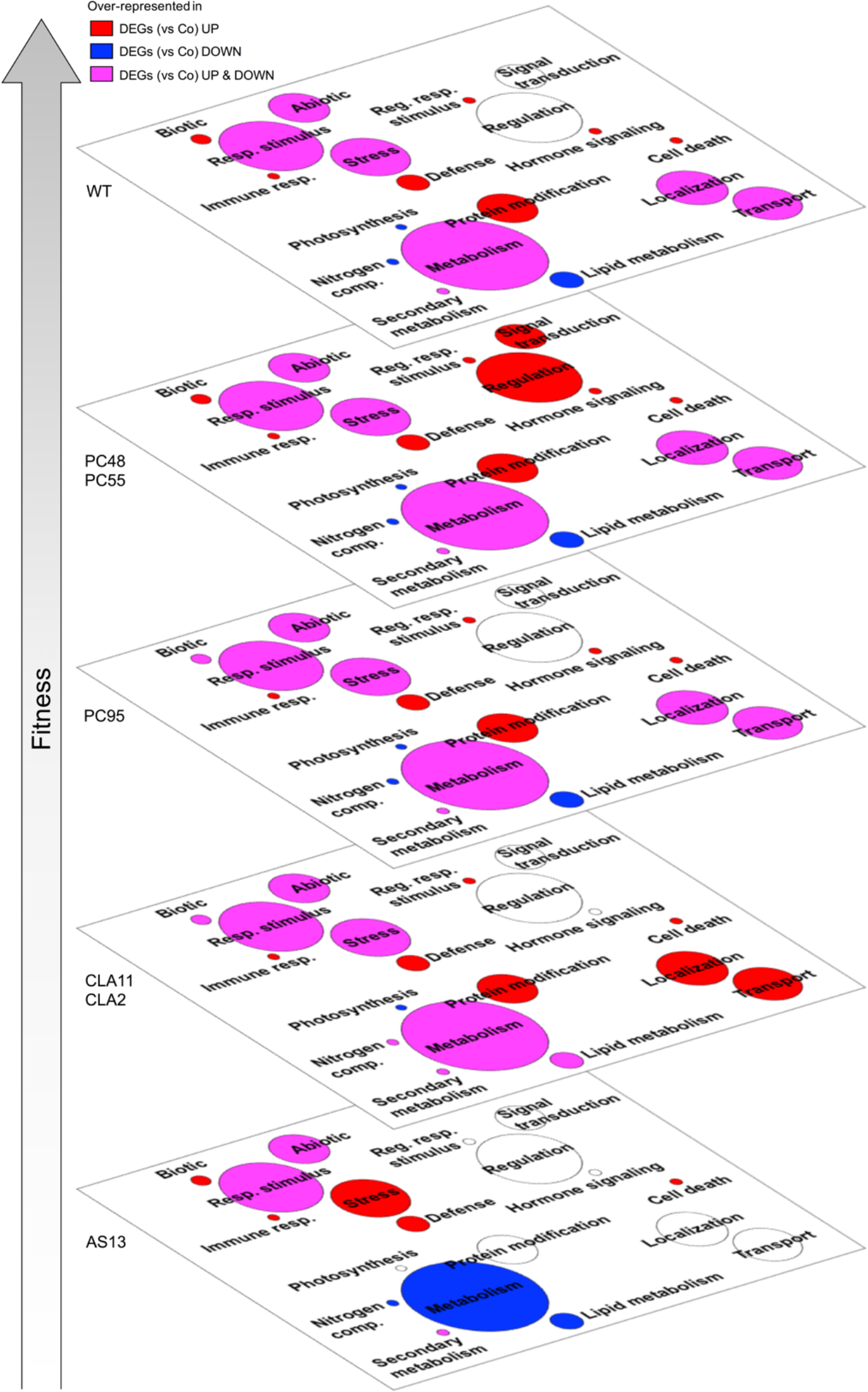
Functional analysis associated to differential gene expression. Artwork of meaningful biological processes (in a plane), where those that are over-represented in the two lists of DEGs for each TEV genotype (up-and down-regulated), either in one of them or in both, are colored. Enrichments evaluated by Fisher’s exact tests with FDR, adjusted *P* < 0.05. The different planes are organized according to viral fitness. We considered infected *vs.* mock-inoculated control plants.

The second fitness plane corresponds to genotypes PC48 and PC55, both of mild effect and carrying mutation in the *CI* gene. The most remarkable difference between these two genotypes and the rest of genotypes is the significant enrichment in up-expressed DEGs related to *signal transduction* (GO:0007165) and *regulation of gene expression* (GO:0010468). Drifting down in the virus’ fitness scale, the next plane in Fig. 5 corresponds to genotype PC95, which shows a similar distribution of GOs than the WT except a loss of enrichment in up-expressed DEGs in the *regulation of response to biotic stress* category. Next plane corresponds to genotypes CLA2 and CLA11 of moderate fitness and carrying point mutations in the *HC-Pro* gene. Plants infected with these two viral genotypes differ from plants infected with the WT virus in three main functional categories: the loss of significant enrichment in the *hormone-mediated signaling* category, and a significant enrichment in up-expressed DEGs into the *localization* and *transport* categories. Finally, the bottom plane in the fitness scale corresponds to genotype AS13 (Fig. 5), which has very low fitness and induces no symptoms or very mild symptoms. These differences in fitness and severity of symptoms have a direct translate into the enrichment of the different functional categories. Compared to WT, *metabolic process* is now enriched with down-expressed DEGs whilst unspecific *response to stress* is very much enriched now in up-expressed DEGs, and *response to stimulus*, *protein modification process*, *localization*, and *transport* are not enriched in any particular type of DEGs. Moreover, no significant enrichment in the *hormone-mediated signaling* module was found in plants infected with AS13, likewise it was observed for the other HC-Pro mutant genotypes CLA2 and CLA11.

### Correlation between gene expression and viral fitness

Taken together, the results shown in the previous sections suggest a transcriptomic response of plants to infection varies with the fitness of the virus being inoculated. This observation motivated us to identify genes whose expression significantly correlates with viral fitness; that is, systematic changes in virus fitness are associated with an increase or decrease in the expression level of a particular gene. This is a correlation analysis and as such does not assume a functional dependence between viral fitness and the expression of individual host genes. Yet, it may provide a list of candidate genes to be considered as determinants of viral fitness. We computed a non-parametric Spearman’s correlation coefficient between viral fitness and the normalized degree of expression (*z*-score) for each one of the *N. tabacum* genes previously characterized as DEGs (Fig. 3A). A total of 326 DEGs show a significant positive correlation and 154 DEGs show a significant negative correlation (Fig. 6A; red dots for positively correlated genes, blue dots for negatively correlated ones; data in Supplementary Dataset S4).

Next, we sought to explore which functional categories and molecular functions, if any, were enriched among these two subsets of DEGs. Results are shown in Fig. 6B and functional annotations are all reported in the Supplementary Dataset S4. There are significant differences in the distribution of positively and negatively correlated DEGs into different functional categories (Fig. 6B, left column: *χ*^2^ = 29.225, 6 d.f., *P* < 0.001), being the difference moderate in magnitude (Cramér’s *V* = 0.304). Amongst DEGs whose expression is positively correlated with viral fitness, *biological regulation* (GO:0065008) and *developmental processes* (GO:0032502) are strongly enriched compared with the negatively correlated DEGs. By contrast, negatively correlated DEGs disproportionally contribute more than positively correlated ones to the categories *response to stimulus*, *localization*, *metabolic processes*, and *cell death*. Focusing in molecular functions (Fig. 6B, right column), a significant difference also exists among DEGs whose expression is positively-and negatively correlated with viral fitness (*χ*^2^ = 36.720, 6 d.f., *P* < 0.001), being the magnitude of the difference in the moderate to large magnitude range (Cramér’s *V* = 0.341). On the one hand, among positively correlated DEGs, *nucleic acid binding* (GO:003676) shows the largest departure from negatively correlated ones. On the other hand, *catalytic activity* (GO:003824) *and transporter activity* (GO:0005215) are the two molecular functions that appear as enriched among negatively correlated DEGs. Together, these results suggest that positively correlated DEGs play a role in the transcriptional regulation of host defenses. By contrast, DEGs with negative correlation between expression and TEV fitness participate more in catalytic and transport activities than genes with positive correlation, suggesting a redirection of resources by the host to face the viral infection, which is not independent of viral fitness.

**Figure 6.**
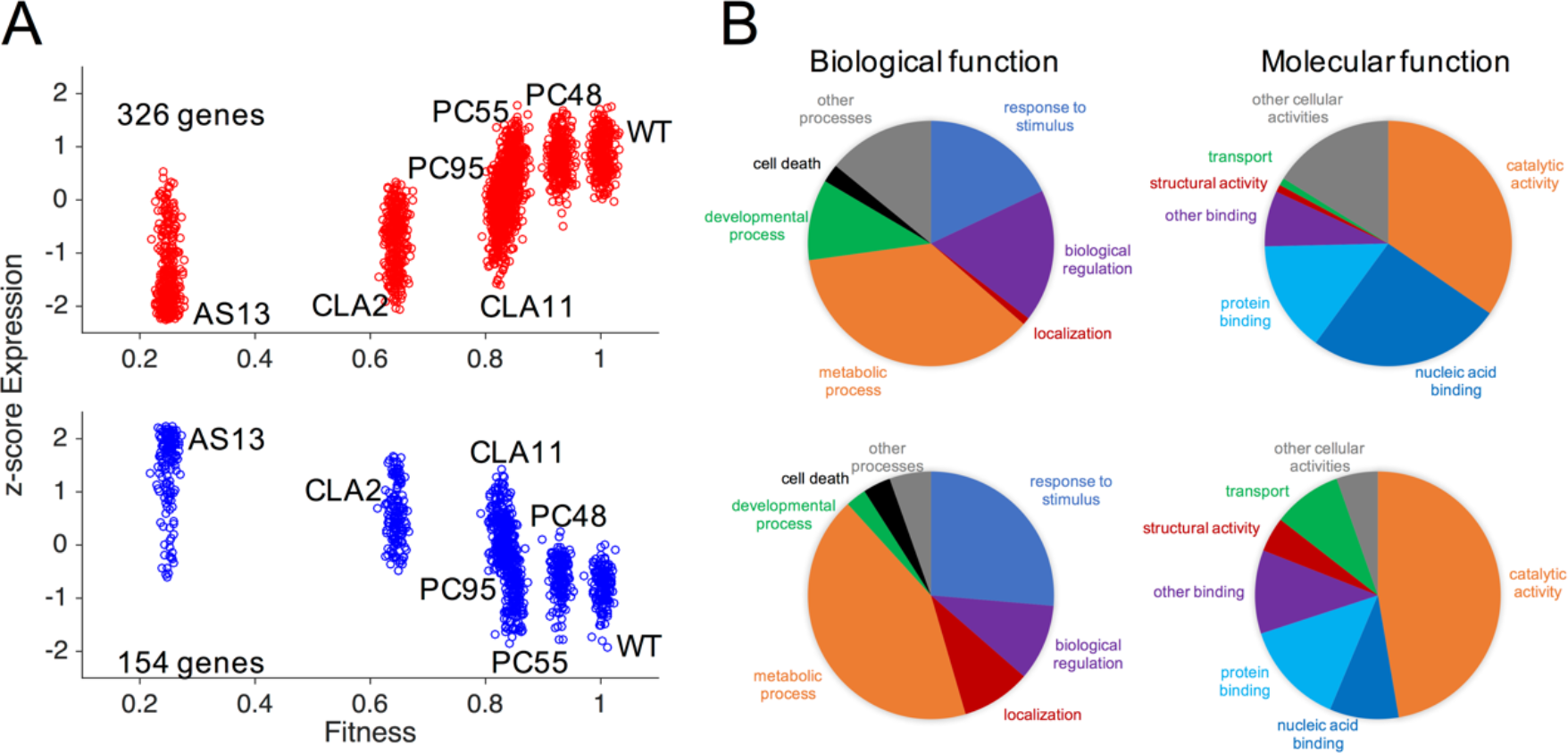
Association between TEV fitness and plant gene expression. **(A**) Correlation plots between host gene expression and viral fitness for those genes that significantly vary across all viral infections (ANOVA with FDR, adjusted *P* < 0.05), and that exhibit a significant positive (upper panel; red dots) or negative (lower panel; blue dots) trend (Spearman’s correlation test, *P* < 0.05). Expression data represented as *z*-scores. **(B)** Pie charts of biological and molecular functions. On the top, for genes whose expression increases with TEV fitness [red dots in panel (A)]; on the bottom, for genes whose expression decreases with fitness [blue dots in panel (A)].

### Validation of microarray data by RT-qPCR for nine representative genes

Normalized expression data used in Fig. 6 were estimated from changes in spot intensity in the *N. tabacum* Gene Expression 4×44K Microarrays (Agilent). To validate these results with an alternative method, RT-qPCR, we selected four positively correlated and five negatively correlated DEGs that cover the entire range of observed significant Spearman’s correlation coefficients (Supplementary Dataset S4). They represent different biological functions and are expressed at different developmental stages and under different environmental situations, so they are not a biased sample (see below). The four positively correlated DEGs selected were (ordered according to the observed *r*_*S*_ values): *dicer-like 2* gene (*DCL2*; *r*_*S*_ = 0.893), the gene encoding for the VQ motif-containing protein 29 (*VQ29*; *r*_*S*_ = 0.893), the gene encoding for the GAST1 protein homolog 1 (*GASA1*; *r*_*S*_ = 0.857), and a gene encoding for a member of the lipase/lipoxygenase PLAT/LH2 family (*PLAT1*; *r*_*S*_ = 0.786). *DCL2* is involved in defense response to viruses, maintenance of DNA methylation and production of ta-siRNAs involved in RNA interference^42^. *VQ29* is a negative transcriptional regulator of light-mediated inhibition of hypocotyl elongation that likely promotes the transcriptional activation of *phytochrome interacting factor 1* (*PIF1*) during early seedling development, participates in the jasmonic acid-mediated (JA) plant basal defense and the VQ proteins interact with WRKY transcription factors^43^. *GASA1* encodes for a gibberellin-and brassinosteroid-regulated protein possibly involved in cell elongation^44^, also reported to be involved in resistance to abiotic stress through ROS signaling^45^. *PLAT1* encodes for a lipase/lipoxygenase that promotes abiotic stress tolerance^46^ and is a positive regulator of plant growth and regulates the abiotic-biotic stress cross-talk. The negatively correlated DEGs selected for validation are the *adenosine kinase 2* gene (*ADK2*; *r*_*S*_ = −0.857), the gene encoding for the small 3B chain of the Rubisco (*RBCS3B*; *r*_*S*_ = −0.857), the *AGAMOUS-like 20* gene (*AGL20*; *r_S_* = −0.786), the *factor of DNA methylation 1* gene, (*FDM1*; *r*_*S*_ = −0.786), and the *granule-bound starch synthase 1* gene (*GBSS1*; *r*_*S*_ = −0.821). *ADK2* encodes for an adenosine kinase involved in adenosine metabolism, including the homeostasis of cytokinines^47^, controls methyl cycle flux in a SAM-dependent manner and plays a role in RNA silencing by methylation. *RBCS3B* is involved in carbon fixation during photosynthesis and in yielding sufficient Rubisco content^48^. *AGL20* is a DNA-binding MADS-box transcription activator modulating the expression of homeotic genes involved in flower development and maintenance of inflorescence meristem identity, transitions between vegetative stages of plant development and in tolerance to cold^49^. *FDM1* is an SGS3-like protein that acts in RNA-directed DNA methylation participating in the RNA silencing defense pathway^50^. *GBSS1* is involved in glucan biosynthesis and responsible of amylase synthesis essential for plant growth and other developmental processes^51^.

Details on the primers used for amplifications, the size of the amplicons, the GenBank identification IDs, and RT-qPCR *C*_*T*_ values for the nine DEGs and the corresponding internal reference genes from the same samples (the ribosomal protein L25 and the translation elongation factor EF1α; ref. 52) are all reported in Supplementary Dataset S5 (see Methods for details). Relative expression data were calculated using the DD*C*_*T*_ method normalized by each one of the two reference genes and then averaged. Finally, to make expression data by both methods (microarray readings and RT-qPCR) readily comparable, they were both transformed into *z*-scores. The results are shown in Fig. 7. Fig. 7A shows the comparison of the two expression measures for the four DEGs with positive correlation with TEV fitness and Fig. 7B for the five DEGs with negative correlation with TEV fitness. Two different plots are presented for each gene. In all cases, the left plot illustrates the relationship between the expression *z*-scores obtained with the microarray method (*x*-axis) and with RT-qPCR method (*y*-axis) for each one of the seven TEV genotypes; the solid lines indicate the best linear fitting between these two datasets. In this representation, a regression line of slope 1 is expected if both quantification methods provide identical *z*-scores. In all nine cases, both expression *z*-scores are highly and significantly correlated; Pearson’s *r* values ranged from 0.696 (*VQ29*) to 0.970 (*GASA1*) (in all cases 5 df, 1-tailed *P* ≤ 0.041). If a more stringent Holm-Bonferroni correction of the overall significance level is taken, then *VQ29* would not remain significant.

**Figure 7.**
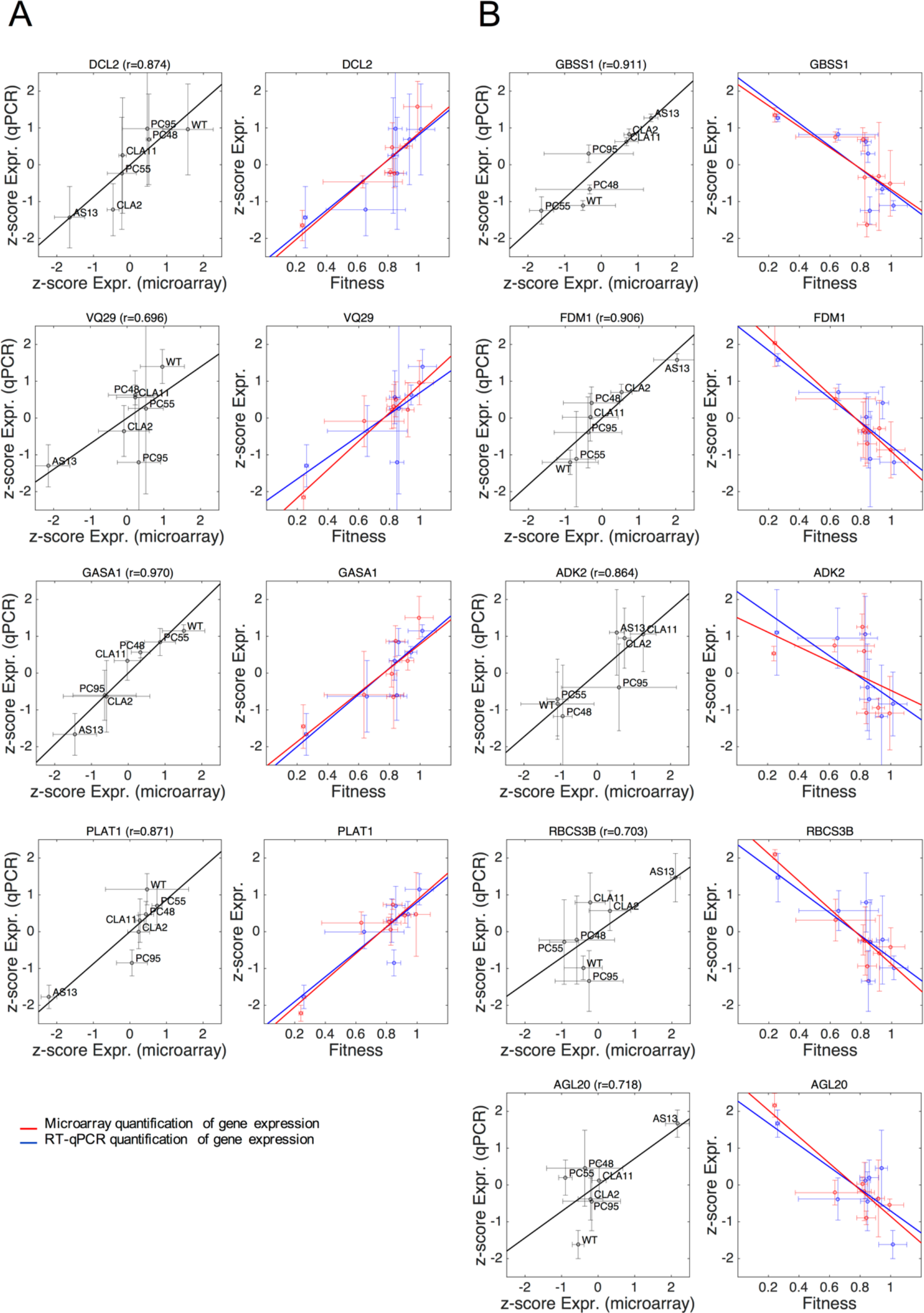
Validation of the transcriptomic data by RT-qPCR for a set of selected genes. For each gene, two different scatter plots are presented. The left panel shows the relationship between the normalized (*z*-score) expression levels measured by transcriptomics with microarrays (*x*-axis) and by RT-qPCR (*y*-axis). The solid line represents the null hypothesis of equal expression values (*i.e*., a perfect match between both quantifications). The Pearson’s correlation coefficient (*r*) is shown on the top of each panel. The right panel shows the correlation plots between gene expression (in red measured by microarray and in blue by RT-qPCR) and viral fitness. The solid lines represent linear models; the closer the slopes of both lines, the more similarity between microarray and RT-qPCR expression data. **(A)** Genes whose expression increases with viral fitness (cases from Fig. 6A, red dots). **(B)** Genes whose expression decreases with viral fitness (cases from Fig. 6B, blue dots). Bidimensional error bars represent ±1 SD.

For each gene, the right plot shows both expression *z*-scores as a function of TEV fitness; solid lines represent the best linear fitting between normalized expressions and TEV fitness. In this representation, the more overlap between the two regression lines, the better the agreement between both quantitative methods. In this representation, *VQ29* and *ADK2* show the largest departure between both regression lines, thought even in these extreme cases, the difference was not large enough as to be significant in a non-parametric Wilcoxon’s signed ranks test (*P* ≥ 0.499 in all nine cases) or in a Student’s *t*-test for the comparison of regression coefficients (*P* ≥ 0.285). Thus, we conclude that, at least for the sample of genes here analyzed, the observed correlations between host’s gene expression and viral fitness are consistent for both experimental methods used to evaluate the levels of gene expression.

### A model of TEV infection incorporating the effect of viral fitness

We further delineated a picture of virus-plant interaction reflected in precise alterations of transcriptomic profiles and regulatory networks. Plant-virus interactions result from the confrontation of two players with opposed strategies and interests. From the plant perspective, activation of basal defenses, immunity, hormone-regulated pathways, and RNA-silencing (some of which are not virus-specific) will result in an immediate benefit to control virus replication and spread. We found that there are plant’s defense responses that are expressed upon infection regardless the fitness of the virus, and defense responses induced progressively as viral fitness increases. Consistent with the first mode, we observed the activation of the genes *EDS1* and *PAD4*, components of R gene-mediated disease resistance with homology to lipases, in every infection studied in this work (Fig. 8A). These are master regulators of plant defenses that connect pathogen signals with salicylic acid (SA) signaling^53^. SA is involved in resistance to a broad spectrum of pathogens, and in particular viruses^54,55^. Consistent with the second mode, we observed the activation of many genes involved in defenses in an extent that is proportional to TEV fitness (Fig. 8A). For example, the *DCL2* and *AGO1* genes-key for the RNA silencing response-, genes modulating resistance to pathogens such as the subtilisin-like protease (*SBT1.9*), or genes expressing proteins involved in hormone-regulated defenses such as *GASA1* and *VQ29*, brassinosteroids (*e.g.*, brassinosteroid enhanced expression 2 *BEE2*, brassionosteroid-signaling kinases 2 and 7, brassinosteroid *BAK1 BRI1*-associated receptor kinase), ethylene response factors (*e.g*., *ERF1B*, *ERF71*, or cytokine response factor 1 *CRF1*), and members of abscisic acid (ABA) perception pathway (*e.g*., *PYL4-RCAR10*, a regulatory component of the ABA receptor family). Likewise, genes involved in methylation-mediated stress responses, such *ADK2*, *FDM1* or the methionine adenosyltransferase *MAT3* reduce their expression as virus replication is more efficient, thus resulting in less methylation and increased expression of genes that participate in apoptosis and post-transcriptional gene silencing^47^. In this way, the overexpression of genes that modulate histone acetylation or chromatin organization, such as the histone acetyltransferase *HAC1* and the chromatin remodeling factor R17 (*CHR17*) would regulate differentiation, apoptosis, transcriptional activation or ethylene response just as viral fitness increases.

**Figure 8.**
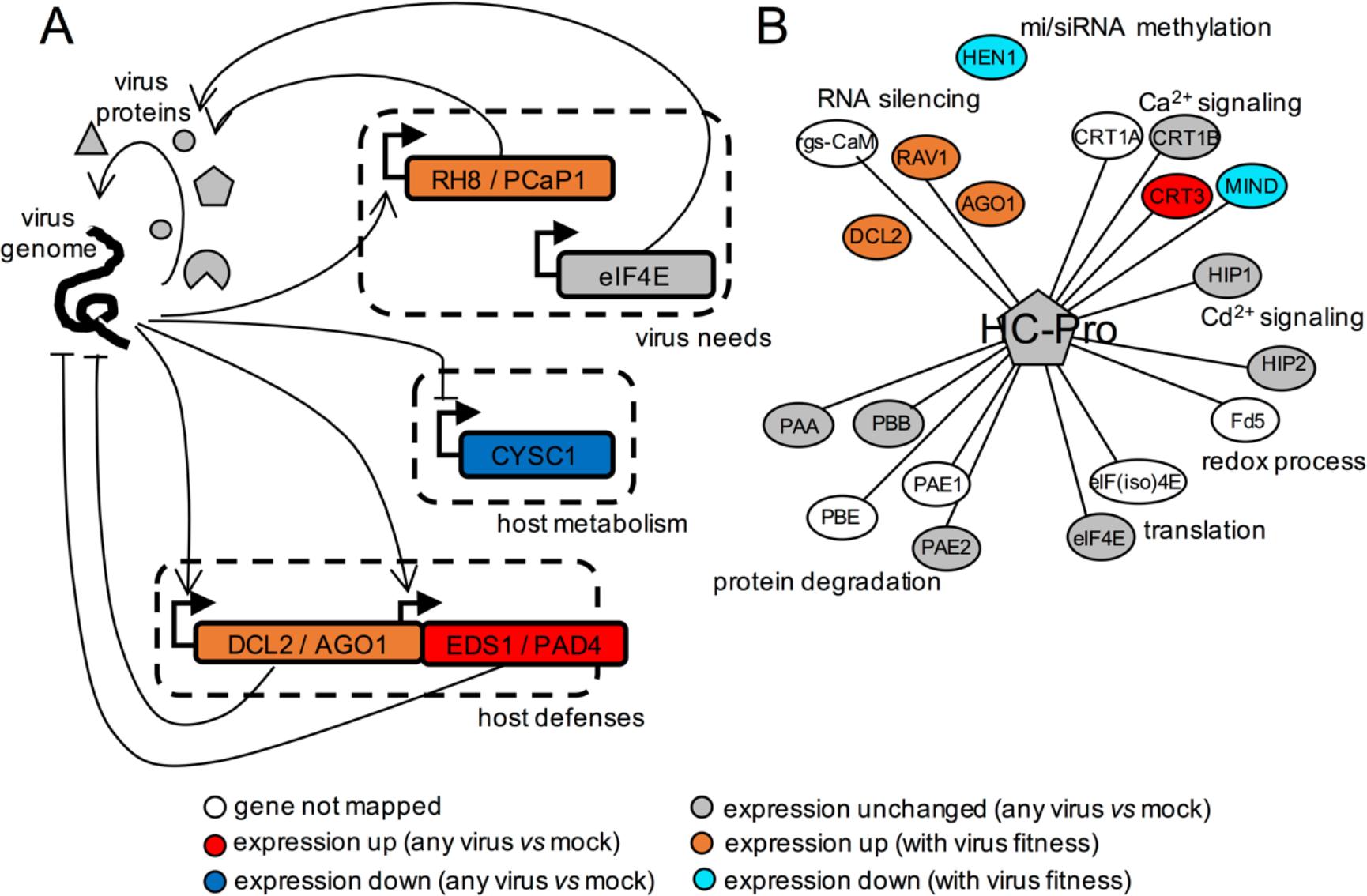
Regulatory model between TEV and plant components. **(A)** Transcriptional regulatory scheme underlying the virus-host interaction. In general, genes participating in host defenses are activated, genes participating in host metabolism are repressed, and genes required by the virus are either activated or unaffected. **(B)** Protein-protein interactome of HC-Pro, the multifunctional protein with VSR activity, contextualizing gene expression data over multiple TEV infections. Different biological processes are indicated.

However, these activations have a cost, mainly in terms of resources that can be invested into secondary metabolism and development. Consistent with this idea is the fact that many genes participating in metabolic processes (*e.g.*, *CYSC1*, a cysteine synthase) are highly repressed upon infection (Fig. 8A). There are also central genes for the plant metabolism whose repression correlates with viral fitness, such as *GBSS1*, photosystem components or assembly factors (*e.g*., *LHCB1.3* and *HCF136*), Rubisco subunits and ATPases, catalases, transketolases, nucleotide and phosphate transporters, synthases involved in flavonoid, isoprenoid, ascorbate, or tryptophan biosynthesis, and GAPDH (Fig. 7).

Diverting host cell resources and reprograming the metabolic machinery to support RNA metabolism and ATP production is a general strategy both of plant^56^ and animal viruses^57,58^. TEV achieves this reprogramming by altering the expression of a series of genes in its own benefit. For example, we found the expression of genes involved in actin cytoskeleton organization such as *ADF4* and *PFN3* to be negatively correlated with TEV fitness. The profilin PNF3 is a actin-binding protein and ADF4 participates in the depolymerization of actin filaments that results from microbial-associated molecular patterns being recognized by the corresponding pattern-recognition receptors^59^. Therefore, by downregulating this function, longer and more stable actin filaments are produced that virions use to move around the cell from the ER-associated replication factories to plasmodesmata. Other example is the repression observed for the *UBP1B* gene, a negative regulator of potyvirus translation, that would allow for a more optimal virus accumulation^60^. Genes involved in nonsense-mediated decay (NMD) defenses^61^, such as the ATP-dependent RNA helicase *UPF1*, also show reduction in levels of expression. Other group of proteins that show alteration during viral infection are those involved in protein degradation, via ubiquitination and downstream into the proteasome pathway (*e.g*., ubiquitin-protein ligase 1, *UPL1*; ubiquitin-conjugating enzyme 2, *UBC2*; ubiquitin E2 variant 1B, *MMZ2*; EIN3-binding F box protein 1, *EBF*) or via autophagy (*e.g*., the ATP-driven chaperone CDC48C and the plant autophagy adaptor NBR1). Moreover, TEV activates in a fitness-dependent manner the expression of genes *RH8*, an RNA helicase, and *PCaP1*, a membrane-associated cation-binding protein, also required by potyviruses for cell-to-cell movement^62^ (Fig. 8A), and of a diversity of transcription factors including global (*e.g*., *GRA2*, *GTE8*), sequence-specific (*e.g*., *SACL1* and *SPL12*), GATA/NAC family members (*e.g*., *GATA1*, *NAC083*, *NAC029*), bZIP G-box finding factors (*e.g*., *GBF1* and *BZIP63*), and involved in homeotic gene expression (*e.g*., *AGL20* and homeobox-1). We also found genes related to genome integrity, (*e.g*., the cohesins *SYN2* and *SMC3*, and the chromatin protein *SPT2*), DNA replication and nucleosome assembly, alternative splicing (*e.g*., *SF1*, an homolog nuclear splicing factor), chromatin transition (*e.g*., *SPT16*, an histone chaperone involved in transcription elongation from RNApolII promoters and regulation of chromatin transitions; or the histone acetyltransferase HAC1, a coactivator of gene transcription with a major role in controlling flowering time and also essential for resistance to bacterial infections), DNA replication and cell division (*e.g*., the mitotic cohesin RAD21 and the cyclin-dependent protein kinase CYCH1). However, not all host factors recruited by the virus present alterations in their expressions. According to our data, the translation initiation factor eIF4E, known to be exploited by TEV for its own translation^63^, was found to be unperturbed (Fig. 8A) whilst eIF3A and eIF4G expression positively correlated with TEV fitness.

In essence, there are genes that are significantly altered (up or down) upon infection irrespective of the ability of the virus to replicate, genes whose expression correlates with this ability (positively or negatively), and genes that remain unaltered. Nevertheless, this picture of virus-plant interaction may be biased by the limited number of viral genotypes analyzed in this work. Three out of six genotypes correspond to *HC-Pro* mutants. As a multifunctional protein, it is not surprising that different fitness levels can be reached by introducing mutations in different functional domains. But, certainly, more mutants should be analyzed in future work to provide a comprehensive picture, avoiding the bias towards certain virus proteins. In addition, we here focused on the transcription regulation, but other networks exist in the cell (*e.g*., metabolism, protein-protein interactions, …), all them interlinked. To provide an insight on these other networks, we constructed the interactome (Fig. 8B) of HC-Pro with the host proteins known to interact with this virus protein^40^. We then contextualized our gene expression data over multiple TEV infections. Many of the cellular functions in which HC-Pro participate (protein degradation, translation, redox processes, and cation signaling) are not regulated transcriptionally upon infection (or regulated marginally). Presumably, the virus exploits these processes in its own benefit (mainly to enhance replication and movement within a cell), and the normal expression of the corresponding genes is sufficient for such subversion. By contrast, RNA silencing and methylation are functions involved in defense against pathogens that are quantitatively regulated, as a sort of control strategy exerted by the plant, as long as they are needed, that is, according to viral fitness.

## Discussion

Biological systems and processes can be analyzed and modeled at every scale of complexity. It is expected that components of each level of complexity may contribute to determine the behavior of processes at other levels. The complexity at the molecular level (*i.e.*, the lowest level of biological organization) is astonishing both in terms of possible elements (genes, functional RNAs, proteins, and metabolites) and of interactions among them^24^. Thus, if the components at lower scales of complexity, presumed to be more accessible experimentally, are informative enough about the underlying processes, they result in excellent proxies to rationalize biological systems. In the case of a disease (in plants or animals), the symptoms exhibited by the organism have been traditionally used as macroscopic indicators of what occurred within the organism. This allows diagnosing diseases without the need to performing further inspections. However, symptoms are generally uncoupled from the magnitude of the perturbation at the molecular level in the host (with respect to a healthy state)^24,25^. This is particularly true in the case of a virus-induced disease, a paradigmatic example of a system-wise perturbation^26–30^.

Here, we studied for the first time the use of viral fitness as a mesoscopic indicator of the molecular changes occurring in the host upon infection. After all, the progress of a viral infection depends on the fitness of the virus mutant swarm. Classically, viral fitness has been evaluated by means of parameters describing the absolute growth and accumulation or by competition experiments^3,4^, or even by correlating it with the development of host’s symptoms^4,32^. We focused on the infections exerted by different genotypes of a given virus in the same host. Fitness differences among genotypes are due to several causes. First, they may be a direct consequence of the effect of mutations on viral proteins, perhaps even resulting in altered folding, and thus jeopardizing their functions. Second, in the case of mutations affecting regulatory regions (*e.g*., RNA stems and loops), the effect may be due to altered structural configurations that impede the binding of virus own proteins or of cellular factors. Plenty of examples illustrate the effect of mutations via these two mechanisms^36^. A third, more tantalizing, yet poorly explored possibility is that mutated viral components (*i.e*., RNAs and proteins) may interact in non-optimal ways with the complex network of genetic and biochemical interactions of the cell as a whole. Interacting in non-optimal ways with any of the elements of the host regulatory and biochemical networks may have profound effects in the progression of a successful infection and thereof in viral fitness. In this work, we considered mutations affecting the CI protein (with RNA helicase, ATPase, and membrane activities), the viral replicase NIb, and the HC-Pro protein (VSR, protease, and helper-component during transmission by aphids). Our results point out that fitness, irrespective of what type of mutation is introduced, is a good indicator of how a given mutant reprograms gene expression patterns, into its own benefit or as a consequence of cellular defenses (*e.g.*, Fig. 3C).

Despite the interest of this hypothesis, none of the early studies tackled the relationship between genotype and fitness of the virus and transcriptomic profiles of the host in a systematic manner, but rather focused on comparing two viral genotypes. Evolution experiments simulating the spillover of TEV from its natural host *N. tabacum* into a novel, poorly susceptible one, *Arabidopsis thaliana*, have shown that adaptation of TEV to the novel host (*i.e*., concomitant to large increases in fitness) was associated with a profound change in the way the ancestral and evolved viruses interacted with the plant’s transcriptome, with genes involved in the response to biotic stresses, including signal transduction and innate immunity pathways, being significantly under-expressed in plants infected with the evolved virus than in plants infected with the ancestral one^64^. Further evolution experiments into different ecotypes of *A. thaliana* that differed in their susceptibility to infection illustrated a pattern of adaptive radiation in which viruses were better adapted to their local host ecotype than to any alternative one, but with viruses evolved into more restrictive ecotypes being more generalists that viruses evolved in the more permissive ones^65^. Interestingly, these differences in fitness had a parallelism with differences in the transcriptomic profiles of plants from different ecotypes; the more generalist viruses altering similar genes in every ecotype, while the more specialist viruses altered different genes in different ecotypes^66^. Similarly, *A. thaliana* plants infected either with a mild or a virulent isolate of turnip mosaic potyvirus (TuMV) also showed profound differences in the genes and functional categories altered^67^. In this case, the more virulent strain mainly altered stress responses and transport functions compared to the mild one^67^. In a recent study, the transcriptomic alterations induced in *Nicotiana benthamiana* plants infected either with a wild-type tobacco vein banding mosaic potyvirus (TVBMV) or a genotype deficient in VSR protein HC-Pro were compared^68^. Both transcriptomes differed in many aspects, including repression of photosynthesis-related genes, genes involved in the RNA silencing pathway, the JA signaling pathway, and the auxin signaling transduction^68^.

Altogether, the results reported in this study illustrate the complex interaction between viruses and their native host plants, and how the outcome of this interaction, in terms of viral replication and accumulation, correlates with the expression of host genes (Fig. 8A). Our observation that viral fitness indeed correlates positively or negatively with the expression of certain genes is of particular interest. By simply measuring the fitness of the virus infecting a given host, we may predict the whole genomic profile of the host cell to characterize its state (molecular impact of infection). Moreover, by specifically targeting host genes that are essential for high fitness virus variants but not for milder ones, we may prevent the spreading of the former variants, while still allowing mild variants to replicate and, perhaps, act as attenuated vaccines that enhance the antiviral response of the plant.

## Methods

### Virus genotypes and plant inoculations

The infectious clone pMTEV contains a full copy of the genome of a WT TEV strain isolated from tobacco (Fig. 1A; GenBank accession DQ986288)^69^. Six TEV mutant genotypes were constructed by site-directed mutagenesis starting from template plasmid pMTEV as described in ref. 70 (mutants AS13, CLA2 and CLA11) and ref. 32 (mutants PC48, PC55 and PC95). Table 1 shows the characteristics of the seven genotypes used in the study.

The pMTEV-derived plasmids contain a unique *Bgl*II restriction site. After linearization with *Bgl*II, each plasmid was transcribed with mMESSAGE mMACHINE SP6 kit (Ambion), following the manufacturer’s instructions, to obtain infectious 5’-capped RNAs. Transcripts were precipitated (1.5 volumes of diethyl pyrocarbonate (DEPC)-treated water, 1.5 volumes of 7.5 M LiCl, 50 mM EDTA), collected and resuspended in DEPC-treated water^71^. RNA integrity and quantity were assessed by gel electrophoresis. Also, each transcript was confirmed by sequencing of a ca. 800-bp fragment circumventing the mutation site as described elsewhere^72^. In short, reverse transcription (RT) was performed using M-MuLV reverse transcriptase (Thermo Scientific) and a reverse primer outside the region of interest to be PCR-amplified for sequencing. PCR was then performed with Phusion DNA polymerase (Thermo Scientific) and appropriate sets of primers for each transcript. Sequencing was performed at IBMCP Sequencing Service. Templates were labelled with Big Dyes v3.1 and resolved in an ABI 3130 XL machine (Life Technologies).

*N. tabacum* L. cv. Xanthi *NN* plants were used for production of virus particles of each of the seven genotypes (Table 1). The 5’ capped RNA transcripts were mixed with a 1:10 volume of inoculation buffer (0.5 M K_2_HPO_4_, 100 mg/mL Carborundum). Batches of 8-week-old *N. tabacum* plants were inoculated with ~5 μg of RNA of each viral genotype by abrasion of the third true leaf. Inoculations were done in two experimental blocks, the first one including AS13, CLA2, CLA11, PC95, and their controls, and the second one including PC48, PC55 and their corresponding controls. All plants were at similar growth stages. Afterwards, plants were maintained in a Biosafety Level-2 greenhouse at 25 °C under a 16 h light and 8 h dark photoperiod. All infected plants showed symptoms 5 - 8 days-post inoculation (dpi), except the AS13 infected plants, which remained asymptomatic and only showed erratic chlorotic spots. At 8 dpi virus-infected leafs and apexes from each plant were collected individually in plastic bags (after removing the inoculated leaf), with the exception of the AS13 infected plants that were collected at 15 dpi. Next, plant tissue was frozen with liquid N_2_, homogenized using a Mixer Mill MM 400 (Retsch), and aliquoted in 1.5 mL tubes (100 mg each). These aliquots of TEV-infected tissue were stored at −80°C.

### RNA preparations

RNA extraction from 100 mg of fresh tissue per plant was performed using Agilent Plant RNA Isolation Mini Kit (Agilent Technologies) following the manufacturer’s instructions. The concentration of total plant RNA extract was adjusted to 50 ng/μL for each sample. Each RNA sample was re-sequenced again at this stage to ensure the constancy of the genotypes as described above.

Viral loads were measured by absolute real-time RT-quantitative PCR (RT-qPCR), using standard curves^72^. Standard curves were constructed using 10 serial dilutions of the WT TEV genome, synthetized *in vitro* as described above, in total plant RNA obtained from healthy tobacco plants treated like all other plants in the experiment. Quantification amplifications were done in a 20 μL volume, using a GoTaq 1-Step RT-qPCR system (Promega) following the manufacturer’s instructions. The forward (q-TEV-F 5’-TTGGTCTTGATGGCAACGTG-3’) and reverse (q-TEV-R 5’-TGTGCCGTTCAGTGTCTTCCT-3’) primers were chosen to amplify a 71 nt fragment in the 3’ end of TEV genome and would only quantify complete genomes but not partial incomplete amplicons^12^. Amplifications were done using an ABI StepOne Plus Real-time PCR System (Applied Biosystems) and the following cycling conditions: the RT phase consisted of 15 min at 37 °C and 10 min at 95 °C; the PCR phase consisted of 40 cycles of 10 s at 95 °C, 34 s at 60 °C, and 30 s at 72 °C; and the final phase consisted of 15 s at 95 °C, 1 min at 60 °C, and 15 s at 95 °C. Amplifications were performed in a 96-well plate containing the corresponding standard curve. Three technical replicates per infected plant were done. Quantification results were examined using StepOne software version 2.2.2 (Applied Biosystems).

### Fitness evaluations

Total RNA was extracted and virus accumulation quantified by RT-qPCR as described above and detailed previously^72^. Virus accumulation (expressed as genomes/ng of total RNA) was quantified 8 dpi for all genotypes with the exception of AS13, that was quantified 15 dpi. These sampling times assure that viral populations were at a quasi-stationary plateau in *N. tabacum*. These values were then used to compute the fitness of the mutant genotypes relative to that of the WT genotype using the expression *W* = (*R*_*t*_/*R*_0_)^1/*t*^, where *R*_0_ and *R*_*t*_ are the ratios of accumulations estimated for the mutant and WT viruses at inoculation and after *t* days of growth, respectively^32^.

Relative fitness data were fitted to a generalized linear model (GLM) with Normal distribution and an identity link function. Virus genotypes and plant replicates were treated as random factors, with the second factor nested within the first one. Three technical replicates of the RT-qPCR were used to evaluate the magnitude of the error term in the model. This statistical analysis was performed with IBM SPSS version 23.

### Microarray hybridizations

Total RNA was isolated as described above and its integrity assessed using a BioAnalyzer 2100 (Agilent) before and after hybridization. The RNA samples were hybridized onto a genotypic designed *N. tabacum* Gene Expression 4×44K Microarray (AMADID: 021113), which contained 43,803 probes (60-mer oligonucleotides) and was used in a one-color experimental design according to Minimum Information About a Microarray Experiment guidelines^73^. Three biological replicates for each of the six TEV mutant genotypes, four replicates for the WT TEV, plus four mock-inoculated negative control plants were analyzed. Sample RNAs (200 ng) were amplified and labeled with the Low Input Quick Amp Labeling Kit (Agilent). The One-color Spike-in Kit (Agilent) was used to assess the labeling and hybridization efficiencies. Hybridization and slide washing were performed with the Gene Expression Hybridization Kit (Agilent) and Gene Expression Wash Buffers (Agilent) as detailed in the manufacturer’s instructions kits. After washing and drying, slides were scanned with a GenePix 4000B (Axon) microarray scanner, at 5 μm resolution. Image files were extracted with the Feature Extraction software version 9.5.1 (Agilent). Microarray hybridizations were performed at IBMCP Genomics Service.

### Differential gene expression analyses

Inter-array analyses were performed with tools implemented in the BABELOMICS 5 webserver^41^. Firstly, all Agilent files were uploaded together to standardize the expression-related signals using quantile normalization^74^. This process ended in a matrix of normalized expression with genes disposed in rows and samples (TEV genotypes, controls and their replicates) in columns, provided as Supplementary Dataset S1. To compare the expression profiles of two TEV genotypes, the expression level corresponding to mock-inoculated plants (control) was first subtracted.

Secondly, differential expression was carried out by comparing two different samples, including replicates (against mock-inoculated or WT TEV-infected plants), by using the Limma test^75^ with false discovery rate (FDR) according to Benjamini and Hochberg^76^ (adjusted *P* < 0.05). An additional criterion of at least two-fold change in mean expression, *i.e*. |log_2_(fold change)| > 1, was imposed to discard genes presenting minimal increases or decreases. Lists of differentially expressed genes (DEGs), up-or down-regulated, provided in Supplementary Dataset S2.

Thirdly, ANOVA tests were performed to identify genes that vary across all conditions (with FDR as above, adjusted *P* < 0.05). To identify the genes shown in Fig. 3A, the test was done over all samples, including the control. By contrast, to identify the genes shown in Fig. 6A, the test was done over all samples corresponding to infections with distinct TEV genotypes. An additional criterion of significant Spearman’s correlation between mean fitness and mean expression (*P* < 0.05) was imposed in this latter case. Lists of genes whose expressions correlate with viral fitness, either positive or negative, provided in Supplementary Dataset S4.

### Functional analyses from gene lists

The annotation of the individual probes in the Agilent’s tobacco microarray (files 021113_D_AA_20130122.txt and 021113_D_GeneList_20130122.txt provided by Agilent) was updated by BLASTing the oligo sequence file (021113_D_Fasta_20130122.txt) against the most recent version of the *N. tabacum* mRNA database (Ntab-BX_AWOK-SS_Basma.mrna.annot.fna) available at the Sol Genomics Network^77^ (SGN). Sequences not returning a significant BLAST hit were removed from the output. A total of 40,430 annotated probes were generated. In 2,673 cases, more than one probe pointed to the same *N. tabacum* gene (*e.g*., probes A_95_P217927 and A_95_P046006 were both complementary to gene *EB438730*), and in those cases the target appeared twice in the output. Each one of the hits could be associated to an alternatively spliced mature mRNA in the SGN database. We then proceeded to generate the list of *N. tabacum* orthologous genes in the *A. thaliana* genome. To do so, we used BLAST against the TAIR version 10 database of *A. thaliana* cDNAs^78^, with a cutoff *e*-value < 10^−4^. The resulting mapping between *N. tabacum* and *A. thaliana* orthologues is listed in Supplementary Dataset S3.

The determination of the gene ontology (GO) categories over-represented within the lists of DEGs was carried out in the AgriGO webserver^79^ by using the Fisher’s exact test (with FDR adjusted *P* < 0.05 according to Benjamini and Yekutieli^80^ criterion). For the graphical representation, we constructed a plane involving the most relevant categories, depicted as circles with size proportional to the total number of host genes belonging to that category (in log scale). In addition, with the lists of genes whose expression correlates with viral fitness, we calculated the pie charts associated to *i*) biological function and *ii*) molecular function in the PANTHER webserver^81^.

### Validation of gene expression through RT-qPCR

Total RNA was extracted from 100 mg of fresh tissue of plants infected with each one of the seven TEV genotypes as described above. The concentration of total plant RNA was adjusted to 50 ng/μL.

Nine candidate genes were selected to validate the effect of infection with each TEV genotype on their expression. Specific primers were designed for each gene that amplified the matured version of their corresponding mRNAs. Primers were designed using OLIGO Primer Analysis Software version 7 (www.oligo.net).

Gene expression was quantified by RT-qPCR relative to the expression of two housekeeping genes^52^. The first housekeeping gen encodes for the L25 ribosomal protein (GenBank accession L18908). Forward NtL25-F (5’-CCCCTCACCACAGAGTCTGC-3’) and reverse NtL25-R (5’-AAGGGTGTTGTTGTCCTCAATCTT-3’) primers were chosen to amplify a 51 nt long fragment. The second housekeeping gen encodes for the elongation factor 1a (GenBank accession AF120093). For this second gene, forward NtEF1a-F (5’-TGAGATGCACCACGAAGCTC-3’) and reverse NtEF1a-R (5’-CCAACATTGTCACCAGGAAGTG-3’) primers were chosen to produce also a 51 nt long amplicon.

Amplifications were done in 20 μL volume, using GoTaq 1-Step RT-qPCR System (Promega) following the manufacturer’s instructions. The forward and reverse primers for each target gene were chosen to amplify a 68 - 137 nt fragments in the corresponding tobacco mature mRNA. Amplifications were done using an ABI StepOne Plus Real-time PCR System (Applied Biosystems) and the following cycling conditions: the RT phase consisted of 15 min at 37 °C and 10 min at 95 °C; the PCR phase consisted of 40 cycles of 10 s at 95 °C, 34 s at 60 °C, and 30 s at 72 °C; and the final phase consisted of 15 s at 95 °C, 1 min at 60 °C, and 15 s at 95 °C. Amplifications were performed individually for each target gene (with the corresponding set of primers) in a 96-well plate containing three biological replicates and two technical replicates per infected plant. Also, each plate incorporates the two housekeeping genes. Since each plate served for the quantification of a single mature mRNA together with the two housekeeping reference genes, a baseline value of 0.1056, resulting from averaging the threshold baselines of all plates analyzed, was used as default threshold. Quantification results were examined using the StepOne version 2.2.2 software (Applied Biosystems).

Expression data (threshold crossing, *C*_*T*_) and primer sequences are reported in Supplementary Dataset S5.

### Data availability

The microarray data that support the findings of this study have been deposited at NCBI GEO with accession number GSE99838. Processed data are presented in Supplementary Datasets. All other relevant data are available from the corresponding author on request.

## Acknowledgements

We thank Francisca de la Iglesia and Paula Agudo for excellent technical assistance, the EvolSysVir lab members for help, comments and discussions, and Lorena Latorre (IBMCP Genomics Service) and Javier Forment (IBMCP Bioinformatics Service) for their assistance. This research was supported by grants from Spanish Ministry of Economy, Industry and Competitiveness - FEDER (BFU2012-30805 and BFU2015-65037-P to S.F.E. and BFU2015-66894-P to G.R.) and Generalitat Valenciana (PROMETEOII/2014/021).

## Author contributions

S.F.E. designed the study. H.C., S.A., and G.P.B. performed the experiments. G.R., H.C., S.A. and S.F.E. analyzed the data. S.F.E. and G.R. wrote the manuscript.

## Additional information

**Supplementary Information** accompanies this paper at http://www.nature.com/ naturecommunications

**Competing financial interests:** the authors declare no competing financial interests.

---

**Supplementary Dataset S1.** Normalized expression data.

**Supplementary Dataset S2.** Number of altered genes and related functional categories.

**Supplementary Dataset S3.** Mapping between *N. tabacum* and *A. thaliana* orthologues.

**Supplementary Dataset S4.** DEGs showing a significant positive and negative correlations to viral fitness.

**Supplementary Dataset S5.** Expression data for the genes validated by RT-qPCR.

